# Physics-based multiscale mass transport model in drug delivery and tumor microenvironment

**DOI:** 10.1101/031252

**Authors:** Arturas Ziemys, Milos Kojic, Mauro Ferrari

## Abstract

We describe multiscale transport model, which was developed to simulate drug diffusion and convection in tissues and drug vectors. Models rely on material properties and physical laws of transport. Our methods show that drug transport analysis may provide deep insight into mechanisms of pharmacokinetics useful in nanotherapeutics and transport study within tumor microenvironment. Because the method relies on material properties and structures, the approach can help studying phenotypical differences as well.

## I. Drug delivery and tumor microenvironment

The limited understanding of tumor microenvironment and tumor-based pharmacokinetics (PK) - can lead to shortfalls of nanotherapeutics. While systemic PK/PD relationships are scrutinized, tumor microenvironment-based PK remains an underexplored territory hiding answers to improve drug delivery. The reason this is becoming such a significant issue is the advancement of nanoparticle (NP) drug delivery systems, which have the capability to reduce the systemic circulation effects of drug compounds and the concentration effects of these particles in the tumor microenvironment, where nanotherapeutics come to rest. Therefore, oncophysical aspects of transport of nanotherapeutics and its passages through biological barriers are fundamentally important [1, 2].

Our *in vivo* and *in silico* study revealed that phenotypic differences of tumor microenvironment can negate enhanced PK properties of nanotherapeutics [3]. There, capillary collagen of type-IV can be a biophysical marker determining the extravasation of doxorubicin (DOX) by using its liposomal formulation (PLD) as a nanotherapeutic formulation of DOX. It was shown that the 3LL tumors having more collagen than the 4T1 tumors prevent DOX extravasation, in opposite to the 4T1 tumors. Furthermore, our in-depth *in silico* analysis showed that the internalization of drug vectors, for example – drug carrying NP, is critical for the enhanced delivery of payload to tumor microenvironment [4]. But not all answers can be easily reached by *in vivo* experiments and advanced imaging techniques. Computational methods can patch gaps in understanding of the mechanisms. Because drug delivery is closely associated with diffusive and convective transports, physics-based computational method may be of great help in studying mass transport. At the same time, multiscale approach is needed to provide answers for different scales, *e.g.* tumor microenvironment, systemic circulation, cell, sub-cellular, organ or molecular levels.

## II. Illustrative Results of Application of Methods

Further, we provide few examples how our transport model was applied to study drug delivery and tumor microenvironment.

### A. Capillary collagen in tumor microenvironment

Collagen fibers are an important component of capillary walls affecting transport of drugs in healthy or tumor vasculature. We used our *in vivo* and multiscale transport models to study DOX transport and accumulation from plasma in tumor microenvironment. We have compared how collagen structure affects the diffusion flux of a 1 nm molecule of DOX and an 80 nm liposome loaded with DOX (DOX-PLD) in tumor vasculature. We have found that collagen fiber sleeve can control the DOX-PLD permeability, while the transport of DOX was little affected (Fig. 1).

**Figure 1.**
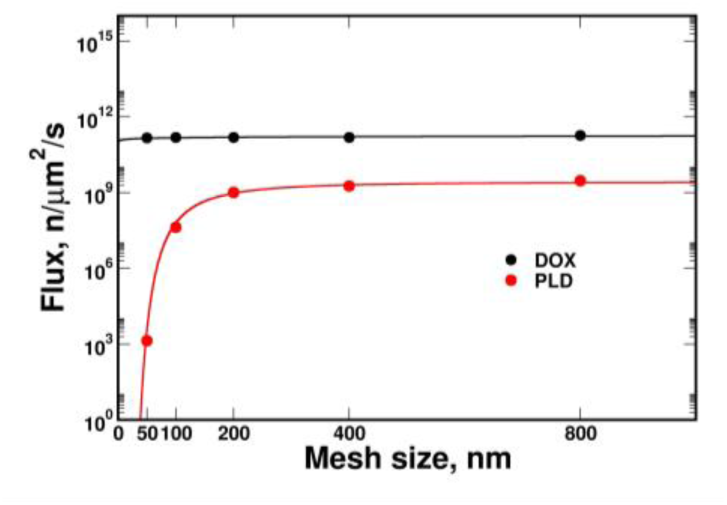
The modeled DOX flux through collagen sleeve of different mesh sizes in two different formulations: free drugs (DOX; *black*) and liposomal formulations (PLD; *red*).

**Figure 2.**
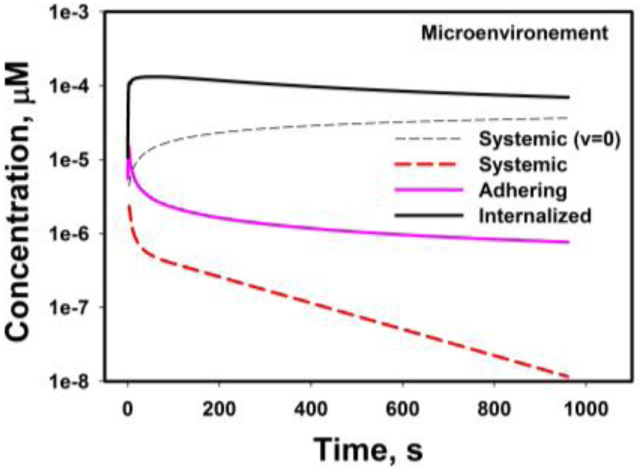
The modeled payload accumulation in capillary microenvironment at different localizations of drug vector and comparison to systemic delivery: systemic delivery and no flow (*black dashed*), systemic with flow (*red dashed*), drug vector adhering on capillary (*magenta solid*), and drug vector internalized into microenvironment (*black solid*).

### B. Drug vector localization for effective drug delivery to microenvironment

We have computationally investigated the interplay of types of physical transport to characterize PK of payload of nanotherapeutics. By comparing payload delivery using systemic circulation and drug vectors to microenvironment, it was found that internalized vectors were the most efficient and showed that the Area under the Curve (AUC) was almost 100 higher than in systemic delivery. The internalization to microenvironment minimizes effects of plasma domain on payload extravasation from nanotherapeutics. The computed results showed that classical PK, which mostly relies on concentration profiles in plasma, sometimes might be inadequate or not sufficient in explaining therapeutic efficacy of nanotherapeutics.

## III. Quick Guide to the Methods (1 Page)

### A. Equations

Here are summarized the fundamentals of the computational model. Considering blood (fluid) and drug vector (solid) as continuous media, the basic equations consist of the balance of linear momentum, 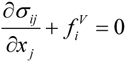 and incompressibility condition 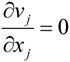, summ on *j*: *j* = 1,2,3 where *σ_ij_* are stresses, 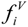 are volumetric forces, and *v_j_* are velocity components of a material point; as indicated, repeated index *j* implies summation *j*=1,2,3 for derivatives with respect to coordinates *x_j_*. Constitutive laws for elastic solid include Young’s modulus *E* and Poisson’s ratio ϑ, and viscosity coefficient μ for fluid. Diffusion is governed by Fick’s law, 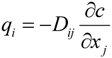, sum on *j*: *j =* 1,2,3 so that the mass balance equation, which includes convection, and can be written as 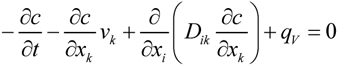, sum on i,*k*: i,*k*=1,2,3 where *q_i_* is mass flux, *v_k_* are Darcy velocities, *c* is concentration, *D_ik_* are diffusion coefficients, and *q_V_* is a source term.

By a common implementation of the principle of virtual power and Galerkin procedure [5], the above balance and continuity equations are transformed into finite element (FE) incremental-iterative algebraic equations for solids, fluids, and diffusion., For diffusion, these FE equations can be written as:

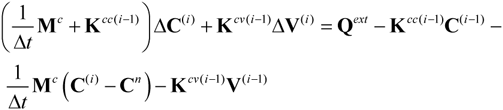

Here, ΔV represent increments of nodal vectors of fluidor Darcy velocity within time step Δ*t*; and Δ**C** is increment of nodal concentration **C**; **Q^ext^** is external mass flux; and the upper right index “*i*” denotes equilibrium iteration number, while the index “n” stands for start of a time step. The above system of equations corresponds to one finite element. This system is further assembled and solved for the nodal quantities for time steps used to calculate evolution of the physical fields of velocity, pressure and concentration; with iterations performed for each time step until convergence is reached [6].

We incorporated concentration and interface effects into FE [7], where diffusivity D is adjusted based on concentration and the proximity to a solid wall, turning D into D(c,w) as D(c,w)=D(c)·S(w). The scaling functions S(w) is derived from MD simulations [7], and incorporates physical effects of the interface.

### B. Type of settings in which these methods are useful

The above methodology can be applied to efficiently study drug mass distribution and its kinetics in tissues and drug vectors, including drug mass release, loading, retention and partitioning. Although transports of physical origin can be modeled directly with some calculated material properties (as diffusivity), the biological transport can also be modelled by using all phenomenological transport coefficients.

## Acknowledgment

The authors acknowledge support from the Houston Methodist Research Institute, Ministry of Education and Science of Serbia (OI 174028, III 41007), City of Kragujevac, and EU grant FP7-ICT-2007 project (224297, ARTreat); the National Institute of Health (U54CA143837 – M.F., K.Y., U54CA151668 - M.F.), the Ernest Cockrell Jr. Distinguished Endowed Chair (M.F.), and the US Department of Defense (W81XWH-09-1-0212) (M.F.).

